# Short tandem repeat mutations regulate gene expression in colorectal cancer

**DOI:** 10.1101/2023.11.29.569189

**Authors:** Max A. Verbiest, Oxana Lundström, Feifei Xia, Michael Baudis, Tugce Bilgin Sonay, Maria Anisimova

## Abstract

Short tandem repeat (STR) mutations are prevalent in colorectal cancer (CRC), especially in tumours with the microsatellite instability (MSI) phenotype. While STR length variations are known to regulate gene expression under physiological conditions, the functional impact of STR mutations in CRC remains unclear. Here, we integrate STR mutation data with clinical information and gene expression levels to study the gene regulatory effects of STR mutations in CRC. We confirm that STR mutability in CRC highly depends on the MSI status, repeat unit size, and repeat length. Furthermore, we present a set of 1244 putative expression STRs (eSTRs) for which the STR length is associated with gene expression levels in CRC tumours. The length of 73 eSTRs is associated with expression levels of cancer-related genes, nine of which are CRC-specific genes. We show that linear models describing eSTR-gene expression relationships allow for predictions of gene expression changes in response to eSTR mutations. Moreover, we found an increased mutability of eSTRs in MSI tumours. Our evidence of gene regulatory roles for eSTRs in CRC highlights a mostly overlooked way through which tumours may modulate their phenotypes. The increased mutability of eSTRs in MSI tumours may be an early indication that eSTR mutations can confer a selective advantage to tumours. Future extensions of our findings into larger cohorts could uncover new STR-based targets in the treatment of cancer.

## INTRODUCTION

Short tandem repeats (STRs) are consecutive repetitions of one to six base pair (bp) motifs. They are estimated to make up around three percent of the human genome (Ellegren, 2004). STRs are a rich source of genetic variation, with individual loci having mutation rates up to 1 *×* 10*^−^*^4^ mutations per generation — 10000 times higher than point mutations (Sun *et al*., 2012). STR mutations are typically the result of DNA polymerase slippage events during replication, where strand misalignment after polymerase detachment results in the insertion or deletion of one or more repeat units.

STR mutations are infamous for their role in neurodegenerative diseases like Hungtington’s disease, which is caused by a large expansion of a protein-coding trinucleotide STR. However, stepwise STR mutations are much more common. These smaller repeat length changes can have functional implications as well, especially when aggregated across the genome (Mitra *et al*. (2021); Verbiest *et al*. (2023)). Apart from inducing frameshift mutations in coding regions, STR length variations have been found to affect DNA methylation (Martin-Trujillo *et al*., 2023) and gene expression (Gymrek *et al*. (2016); Fotsing *et al*. (2019); Shi *et al*. (2023)) under physiological conditions. STR mutations are especially prevalent in tumours of the microsatellite instability (MSI) phenotype. MSI is a type of genome instability where DNA mismatch repair genes are mutated or silenced, causing hypermutability of STRs (Boland and Goel, 2010). MSI is a known subtype in several cancer types. One such cancer type is colorectal cancer (CRC), where around 15-20% of tumours are classified as MSI (Bonneville *et al*., 2017). Given the widespread regulatory effects of STR length variations in healthy tissues, STR mutations may be expected to affect gene expression in CRC as well.

However, the extent to which STR mutations regulate gene expression in tumours remains unclear, despite several large studies quantifying STR mutations in cancer (Hause *et al*. (2016); Bonneville *et al*. (2017); Fujimoto *et al*. (2020)). There has been some evidence suggesting somatic STR mutations may be related to gene expression changes. For example, Bilgin Sonay *et al*. (2015) reported global changes in expression levels between groups of genes with and without STR mutations in promoter regions in CRC tumours. Others have linked individual STR mutations to expression changes of single genes in cancer (Kim *et al*. (2013); Maruvka *et al*. (2017)). Even with these findings, however, what is still lacking is an in-depth analysis of which STR loci regulate the expression of which genes through variations in their allele lengths in tumours.

Here we report a systematic study of stepwise STR mutations in CRC, and present evidence of gene expression changes mediated by somatic mutations affecting STR lengths. First, we generated a new reference panel of STR loci for all human protein-coding genes. Then, we used this panel as a basis for STR genotyping and mutation calling in whole-exome sequencing (WES) data from The Cancer Genome Atlas (TCGA) (The Cancer Genome Atlas Network, 2012). Through this analysis, we could confirm many previously reported factors that influence STR variability. Moreover, we identified a set of expression STRs (eSTRs) for which the allele lengths were associated with expression levels of nearby genes in CRC tumours. We could demonstrate that this eSTR panel allows for predictions of gene expression changes in response to somatic STR mutations in patient-matched samples. Finally, we observed an increase in mutability of eSTRs in MSI tumours, suggesting that eSTR mutations may be under positive selection in cancer under some circumstances.

## METHODS

### Generating a novel STR panel for human protein-coding genes

STRs were annotated across all protein-coding genes (introns, exons, 5kb upstream) of the GRCh38 human reference genome. Gene coordinates were based on GENCODE version 22 (the version used in the Genomic Data Commons (GDC) data release 31.0). STRs were defined as tandem repeats where the unit size was one to six bp long, and the allele length was at least nine, four, four, three, three, and three units for mono- to hexanucleotide repeats, respectively. These thresholds were chosen by taking the lowest allele length for each unit size reported in Lai and Sun (2003), Willems *et al*. (2014), and Mousavi *et al*. (2019). PHOBOS (Mayer *et al*., 2010), XSTREAM (Newman and Cooper, 2007), and TRF (Benson, 1999) were used to detect STRs in the gene sequences. These algorithms also return imperfect STRs, which can have mismatches and inconsistent unit lengths due to point mutations, insertions, and deletions in individual units. Annotating such repeats in biological sequences is not trivial, and the different heuristics used by the various detection tools can lead to competing annotations of the same STR region. To address this, the outputs of the three repeat detection tools were integrated and harmonised using the dedicated statistical framework implemented in TRAL (Schaper *et al*. (2015); Delucchi *et al*. (2021)). TRAL was used to calculate unit divergence metrics and a *P*-value for every STR detected by the different detection algorithms. With these metrics, the best STR representation for overlapping annotations was selected, yielding a non-redundant STR set. Next, circular-profile hidden Markov models (cpHMMs) were generated for this set of non-redundant STRs. The cpHMMs were used to re-annotate the STRs in the genes using HMMER (Eddy, 2011), allowing for more sensitive detection of repeat boundaries. The heuristic nature of the repeat detection algorithms combined with TRAL’s stringent filtering approach meant that some STRs may have been missed — particularly in low-complexity regions. To address this, the panel was supplemented with STRs detected by PERF, an exhaustive algorithm for detecting perfect STRs (Avvaru *et al*., 2018).

### Genotyping STRs from short-read sequencing data

The STR panel was genotyped in aligned WES data from TCGA COAD and READ cohorts using GangSTR (Mousavi *et al*., 2019). In total, alignments for 377 primary tumour samples, 161 blood derived normal samples, and 28 solid tissue normal samples were downloaded from the GDC knowledge base data release 31.0 (the most recent version at the time of analysis). The STR panel was filtered to ensure high-confidence STR length calls before genotyping. Where possible, imperfect TRAL repeats with inconsistent unit sizes were split into smaller STRs with consistent unit size, while still allowing for mismatches between units. Additionally, STRs located in segmental duplications, on non-autosomal chromosomes, or within 50bp of another STR were removed. This filtered STR panel was used to run GangSTR on the alignments with additional flags --output-readinfo --nonuniform --include-ggl --verbose. Subsequently, low confidence calls were removed using DumpSTR (Mousavi *et al*., 2021) with the following settings: --gangstr-min-call-DP 20 --gangstr-max-call-DP 1000 --gangstr-filter-spanbound-only --gangstr-filter-badCI --zip --drop-filtered. Samples with a low number of calls (*<* 10000 STR loci) were discarded. Finally, somatic copy number variant (CNV) segment data for tumor samples were retrieved from the Progenetix database (Huang *et al*., 2021) and labelled using the labelSeg R package (Zhao and Baudis, 2023). Out of the 4177969 STR loci observed across tumor samples, 616166 (14.75%) overlapped a CNV event. All such STR calls were removed.

Overall, this yielded STR genotypes for 350 primary tumours and 159 healthy samples (blood- or solid-tissue derived). Out of the 350 primary tumours, 286 were annotated as microsatellite stable (MSS) or microsatellite-instability low (MSI-L) by TCGA. As recommended by Kim *et al*. (2013), these two subsets were merged and will be referred to collectively as MSS samples. The other 64 primary tumours were labelled microsatellite instability high (MSI-H) by TCGA, which will be referred to as MSI here.

### Calling somatic STR mutations in CRC tumours

For 145 primary tumour samples (120 MSS, 25 MSI), there was a patient-matched blood derived normal sample or solid tissue normal sample to use for STR mutation calling. There were 15 patients for whom there was both a blood- and solid tissue-derived sample available. In these cases, the sample with the most STR calls was chosen as the healthy reference. The biallelic STR genotype at each locus was compared between the healthy and tumour sample for all 145 patients. Loci where at least one allele in the tumour sample was different from the healthy sample were considered mutated. Analogous to Mitra *et al*. (2021), we removed loci where the cancer sample appeared to be homozygous for an allele not observed in its healthy reference sample. These calls are likely to stem from heterozygous loci that are spuriously called as homozygous due to allele dropout.

### Identifying and validating expression STRs

The eSTR detection approach used here is based on methods described in Gymrek *et al*. (2016) and Fotsing *et al*. (2019). eSTR detection was performed across 331 primary tumour samples for which WES and gene expression data were available. Transcripts per million (TPM) gene expression values for these samples were retrieved, and genes with a median expression of zero were discarded. Expression values of the remaining genes were quantile normalised to standard normal distributions. STR genotypes were represented as the average length of the alleles at every locus. Only the 15084 STRs that were called in at least 50 patients and for which at least three distinct genotypes were observed were considered for the eSTR analysis. STR-gene pairs were defined by determining in which gene(s) STR loci were located. Since some genomic regions contain more than one gene, this resulted in 16667 STR-gene pairs. For each of these pairs, a linear model was fitted with the mean STR genotype as the independent variable, and the normalised gene expression as the dependent variable. This was done using the ordinary least-squares regression implementation from the statsmodels Python library version 0.13.2 (Seabold and Perktold, 2010). A T-test was performed for each model to determine whether the association between the STR length and normalised expression was significantly different from zero. The Benjamini-Hochberg procedure was used to control the false discovery rate across all STR-gene pairs at *α* = 0.05. STRs significantly associated to the expression of a gene after correcting for multiple testing were considered putative eSTRs.

To validate the putative eSTRs, a validation set was defined which consisted of 16 patients that had WES and gene expression data available for both a primary tumour sample and a solid tissue normal sample. The primary tumour samples for these patients were not part of the 311 primary tumour samples used for the eSTR discovery. TPM gene expression values for the validation samples were quantile normalised to the same quantiles that had been generated during the normalisation of the discovery samples. A set of 1493 somatic mutations at putative eSTRs was detected using the approach described above for the general STR mutation calling. These eSTR mutations were used to assess whether the linear eSTR models could predict gene expression changes in response to somatic eSTR mutations.

### Comparing the mutability of eSTRs and non-eSTRs

For the 15084 loci that were included in the eSTR analysis, the mutability of eSTRs and non-eSTRs was compared. This analysis was performed for MSS and MSI patients separately. Since STR mutability depended strongly on the repeat unit and allele length (Results), STRs were grouped into ’repeat types’ based on their unit size and reference allele length. For each repeat type, all observations of eSTRs and non-eSTRs were retrieved from the set of 145 patients for which somatic mutations had been called (see above). Repeat types for which either eSTRs or non-eSTRs were observed fewer than 25 times were discarded. For the remaining repeat types the fraction of non-STRs that was mutated was compared to the fraction of eSTRs that was mutated. The number of repeat types for which a difference in mutability between eSTRs and non-eSTRs was observed was noted. For either eSTRs or non-eSTRs to be considered more mutable for a given repeat type, the difference in the fraction of mutated loci had to be more than 0.05. E.g., if for a particular repeat type 0.41 of eSTRs were mutated versus 0.38 of non-eSTRs, this was not considered to be a difference in mutability. To test whether the observed number of repeat types with a difference in mutability was significantly different from random expectation, this analysis was repeated for 10000 permutations (again for MSS and MSI patients separately). For each permutation, the labels indicating which loci were eSTRs were randomly shuffled, and the fraction of repeat types with a difference in mutability between eSTRs and non-eSTRs was recorded. This resulted in null distributions against which the observed fraction of repeat types with increased mutability of eSTRs or non-eSTRs could be compared using permutation tests.

## RESULTS

### A novel STR panel for human protein-coding genes

To explore STR mutations in CRC, we first annotated STRs in the introns, exons, and promoter sequence of all protein-coding genes in the GRCh38 reference genome (Methods). We discarded STR loci for which genotyping was expected to be inaccurate due to genomic context (Methods), resulting in an STR panel containing 1181838 loci. This panel was genotyped in alignments from the TCGA COAD and READ cohorts (Methods).

Since the TCGA alignments were based on WES data, only a subset of our STR panel could be analysed: 142169 STR loci were genotyped at least once across all samples. The reference coordinates of these genotyped STR loci can be downloaded from: http://webstr.ucsd.edu/ (Lundströ m *et al*., 2023). Most genotyped loci were located in introns, which contained 78.35% of genotyped STRs. Untranslated regions (UTRs), coding sequences (CDS), and promoter regions harboured 9.00%, 8.04% and 4.61% of the genotyped STRs, respectively (Supplementary file 1: Figure S1A). The average reference allele length of STRs was lowest in CDS regions (Mann-Whitney U tests, *P*-value *≪* 0.05 for all pairwise comparisons) (Supplementary file 1: Figure S1B). Additionally, STRs with unit sizes three and six were more likely to occur in CDS regions than STRs of other unit sizes (Supplementary file 1: Figure S1C).

### Repeat characteristics and MSI status affect STR mutability

Using the newly generated STR length calls, we could detect somatic STR mutations in CRC tumours. We did this by comparing the allele lengths of STR loci between patient-matched healthy and tumour samples. This was possible for 145 CRC patients where WES data was available from both a primary tumour and healthy sample (Methods). We considered all instances where the allele lengths at an STR locus in the tumour sample were not identical to those of the patient-matched healthy sample to be STR mutations.

As expected, we found that the proportion of mutated STRs was higher in MSI tumours (on avg. 5.17% of STRs per patient mutated) than in MSS tumours (on avg. 1.40% of STRs per patient mutated) (Mann-Whitney U test, *P*-value *≪* 0.05). Single-step STR mutations were the most common: 55.3% and 35.4% of all STR mutations resulted in a difference of one unit between the healthy and tumour allele in MSS and MSI tumours, respectively (Figure 1A; Supplementary file 1: Figure S2). The average difference in allele length per mutated STR was 2.61 units in MSI tumours. This was significantly larger than for STR mutations in MSS tumours, where the average difference in alle length was 1.93 units (Mann-Whitney U test, *P*-value *≪* 0.05). Furthermore, there was a small but significant correlation between tumour stage and the allele length difference of STR mutations in MSI tumours (Spearman’s *ρ*=0.15, *P*-value *≪* 0.05).

**Figure 1.**
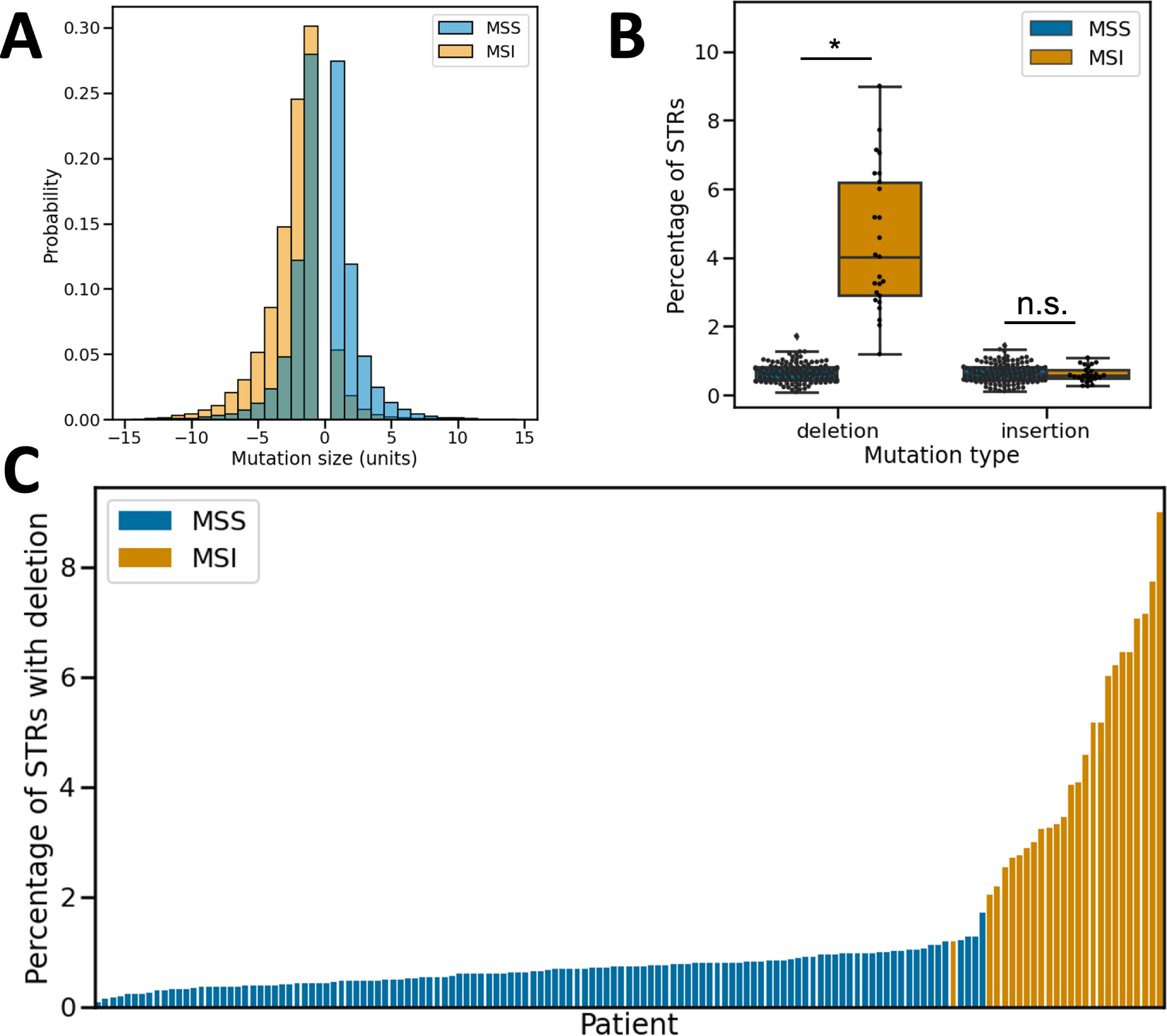
Characteristics of STR mutations in MSS and MSI patients. **A)** Distributions of STR mutation step sizes (in units) for MSS and MSI patients, with negative step sizes indicating deletions in tumours and positive step sizes insertions. Step sizes in the range [-15, 15] are shown. The y-axis displays the probability of an STR mutation being a certain step size. Data from MSS (blue) and MSI (orange) tumours are shown separately as overlapping histograms. The histograms for MSS and MSI data each sum to one. **B)** Boxplots showing insertion and deletion frequencies at STRs in MSS and MSI tumours. Boxes extend from Q1 to Q3, with a line indicating the median value. Significant differences are indicated by asterisks, non-significant differences by n.s. **C)** For every patient for which STR mutations could be called, the STR insertion rate is shown. Patients are ordered along the x-axis based on STR insertion rate, and bars are coloured by MSI status. Abbreviations: MSS, microsatellite stable; MSI, microsatellite instable.

These observations may indicate that at least part of the difference in mutation stepsize between MSS and MSI tumours was caused by multiple mutations occurring per STR in MSI tumours. Mutations in MSS tumours were split evenly between deletions and insertions (23770 vs. 23510, respectively). In MSI tumours, on the other hand, deletions were over ten times more common than insertions (40679 deletions versus 4019 insertions) (Figure 1A). In fact, the higher STR mutation frequency observed in MSI tumours was due solely to deletions, as the fraction of STRs with insertions was not significantly different between MSS and MSI tumours (Mann-Whitney U test, *P*-value 0.42) (Figure 1B). Ranking patients based on the fraction of STRs with deletions yielded a separation into MSS and MSI groups that was highly concordant with TCGA labels (Figure 1C).

The mutability of STR loci was strongly related to their unit size, with repeats consisting of shorter motifs being more variable. This was true for both MSS and MSI patients (Figure 2A). The observed differences in STR mutation frequencies between MSS and MSI patients were caused primarily by mono- and dinucleotide repeats, and to a lesser degree tri- and tetranucleotide repeats. For STRs of these unit sizes, the proportion of mutated loci was significantly higher in MSI patients (Fisher’s exact tests, *P*-values *≪* 0.05). For STRs with unit size five or six, there was no significant difference in mutability between MSS and MSI patients (Fisher’s exact tests, *P*-values *>*0.05) (Figure 2A).

**Figure 2.**
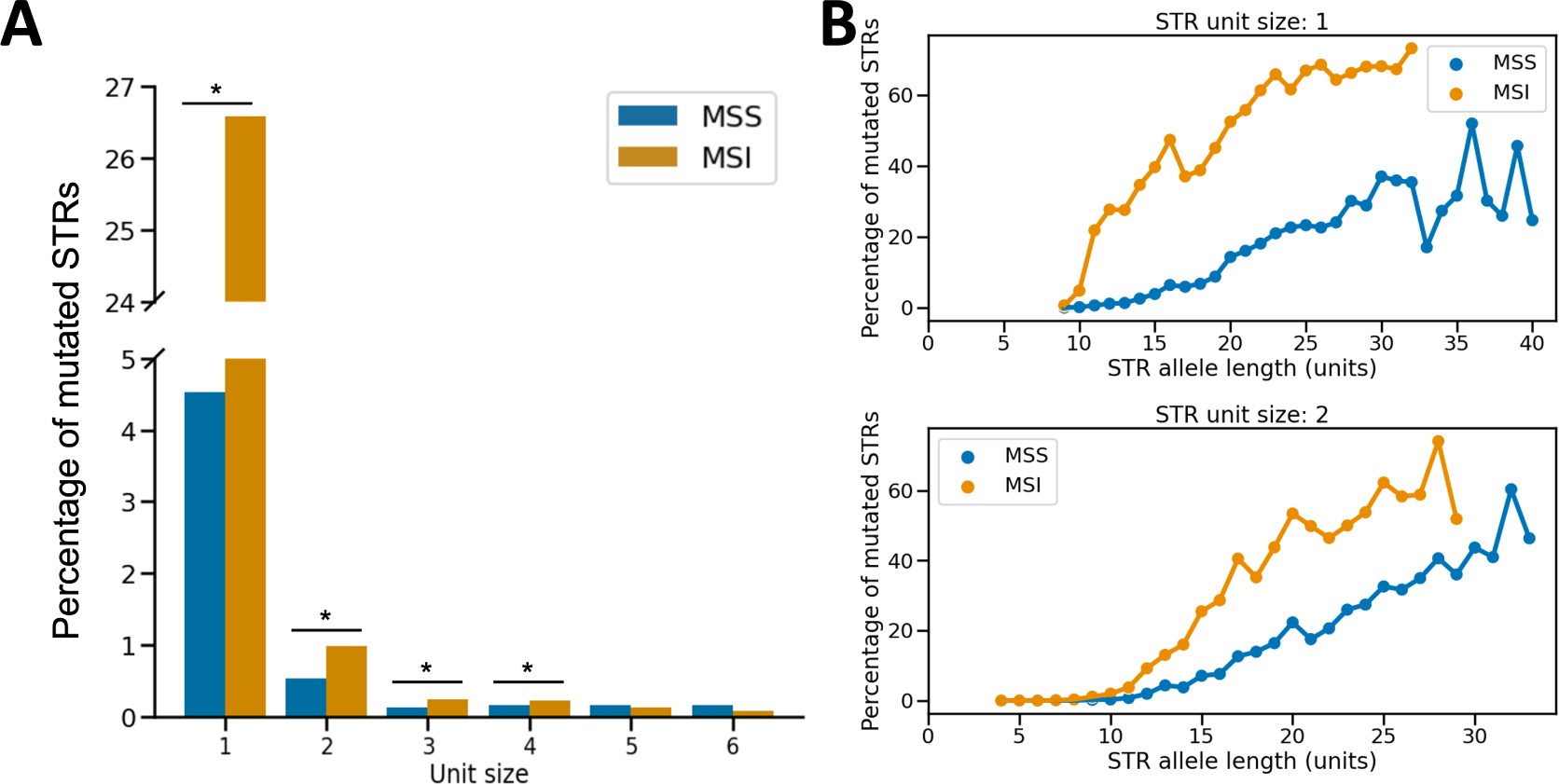
STR unit size and allele length influence mutability in CRC. **A)** STR mutation rates for the different unit sizes. Mutation rates are shown separately for MSS and MSI tumours. Note: the y-axis contains an axis break to accommodate the full range of mutation rates across the different unit sizes. Significant differences in mutation rate between MSS and MSI tumours are indicated with asterisks. **B)** STR mutation rates as a function of unit size and allele length. For unit sizes one and two, mutation frequencies are shown for STR allele lengths for which a comparison between a healthy and tumour sample could be made in at least 50 patients. Results are plotted separately for MSS and MSI tumours. For other unit sizes, see Supplementary file 1: Figure S3. Abbreviations: MSS, microsatellite stable; MSI, microsatellite instable.

Another factor influencing the mutability of STRs was the allele length. This was especially apparent for mono- and dinucleotide repeats, for which a large range of allele lengths could be sampled (Figure 2B, Supplementary file 1: Figure S3). STR mutation frequencies were close to zero for both MSS and MSI patients at STRs where the allele length was low. As the allele length increased, the mutation frequency increased as well. STR mutation frequencies for MSS and MSI tumours diverged as allele lengths increased, with STR mutability in MSI tumours rising more rapidly. The STR mutability became elevated from around allele lengths of ten, ten, ten, seven, six and six units for mono - hexanucleotide repeats, respectively (Supplementary file 1: Figure S3). These observations indicated that our STR panel included the biologically relevant range of allele lengths where mutation rates are elevated.

### Short tandem repeat mutations regulate gene expression in colorectal cancer

Having confirmed that our approach recovers known aspects of STR- and cancer biology, we set out to investigate whether somatic STR mutations affect gene expression in CRC. To identify eSTRs, we fitted linear models between STR allele lengths and gene expression for 15128 STR-gene pairs across 331 primary tumour samples (Methods; Supplementary file 2). We then performed a T-test for each linear model to determine if the slope (*β*) was significantly different from zero. As a negative control, we also performed this analysis with permuted STR genotypes. The *P*-values from the permuted analysis closely followed a uniform distribution, as is expected under the null hypothesis of zero eSTRs (Figure 3A). The *P*-values obtained from the original analysis, on the other hand, were consistently lower than expected under the null model (Figure 3A).

**Figure 3.**
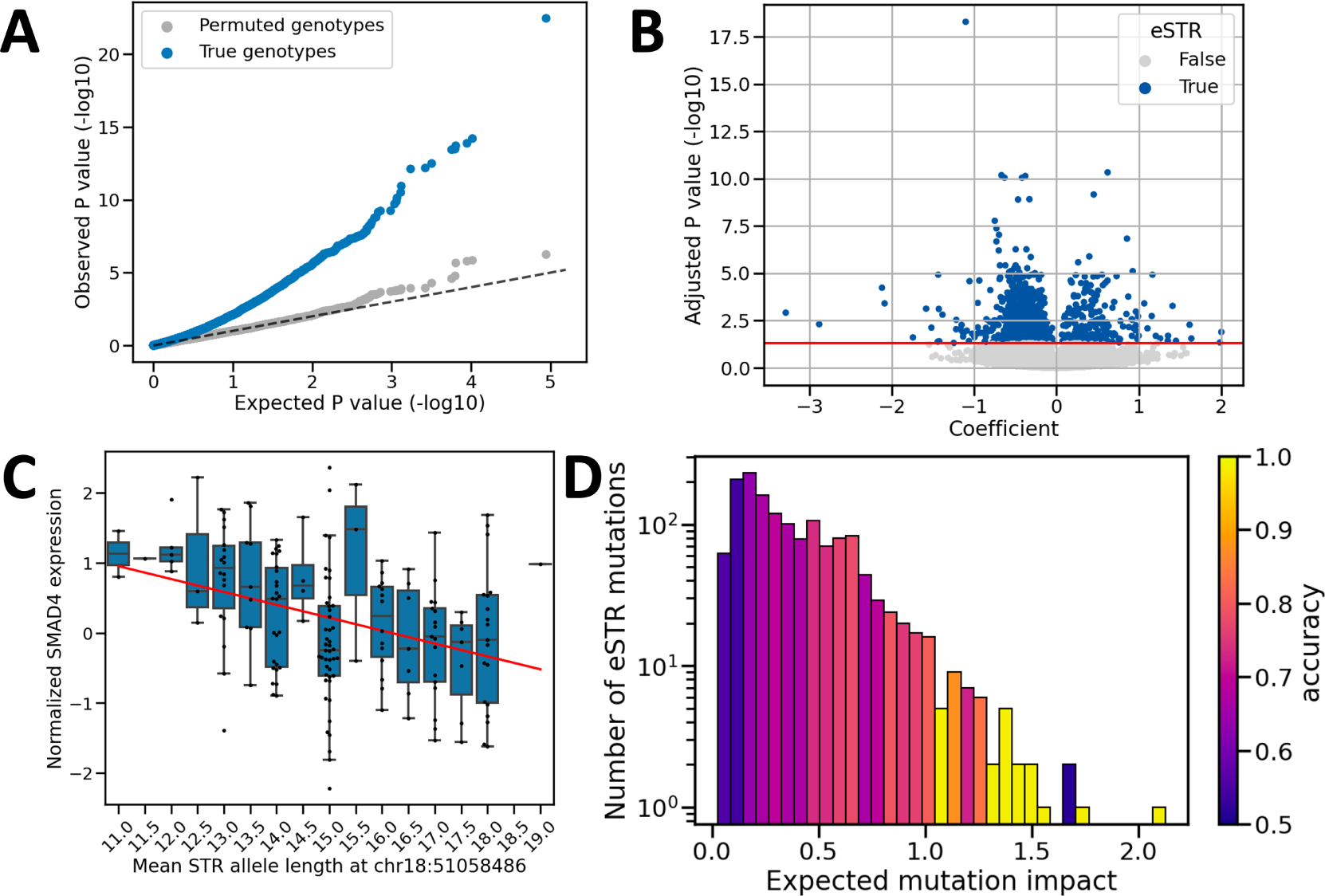
eSTRs detected in colorectal cancer tumours. **A)** Q-Q plot comparing expected versus observed *P*-values obtained from the eSTR analysis (blue) and *P*-values obtained under permutation of STR genotypes (grey). Expected *P*-values were generated under a continuous uniform distribution on the interval [0, 1], representing the null hypothesis of no eSTRs (dashed line). **B)** Significance testing of eSTR-gene expression associations. The coefficients and their *P*-values are plotted for all tested STR-gene pairs. The horizontal red line indicates the significance threshold after controlling the false discovery rate at *α*=0.05 using the Benjamini-Hochberg procedure. Dots are coloured based on significance of the coefficient (grey=not significant, blue=significant). In total, there were 1259 STR-gene pairs with a significant association. **C)** Example of an eSTR. We observed a significant linear relationship between the allele length of a mononucleotide repeat starting at position 51058486 on chromosome 18 and the normalised expression of *SMAD4*. The STR length is shown on the x-axis (mean of two alleles), and the normalised *SMAD4* expression on the y-axis. Every dot represents one tumours sample. Boxplots show the distribution of expression values across tumours at each STR genotype. Boxes extend from Q1 to Q3, with a line indicating the median value. The red line represents the linear model relating STR length to normalised expression. **D)** Histogram showing the expected mutation impact of eSTR mutations. Bars are coloured based on the accuracy of gene expression change predictions obtained using mutations in each bin. eSTR mutations with high expected impact tended to yield higher prediction accuracy. Abbreviations: eSTR, expression short tandem repeat.

A significant relationship between the STR allele length and gene expression was found for 1259 pairs after correcting for multiple testing (T-tests, adjusted *P*-values *<* 0.05) (Figure 3B). We considered these loci to be putative eSTRs. Since some eSTRs were significantly associated with the expression of more than one gene, there were 1244 unique eSTR loci. The vast majority of these loci (973) were mononucleotide repeats located in introns. Among the putative eSTRs, there were 73 for which the allele length was associated with the expression of a cancer-related gene according to COSMIC (Tate *et al*., 2019). Nine of these genes were specifically involved in CRC: *HIF1A*, *KRAS*, *MDM2*, *PTPRT*, *QKI*, *RAD21*, *SMAD2*, *SMAD4*, *TGFBR2* (Supplementary file 1: Figure S4). For ten of these CRC genes the eSTR was located in an intron. The one exception was *TGFBR2*, for which the eSTR was an A/T mononucleotide repeat located in the third exon of the gene. The reference allele length for this STR is 10. Deviations from this number are expected to shift the reading frame and result in premature stop codons. Indeed, *TGFBR2* expression was significantly lower in tumours with non-wild-type allele lengths for this eSTR (two-sided T-test, *P*-value = 0.0018), consistent with nonsense-mediated decay of the transcript.

To isolate eSTR effects from any linked expression quantitative trait loci (eQTL), we investigated the ability of the putative eSTRs to predict gene expression changes in response to somatic mutations. To this end, we leveraged 1493 somatic eSTR mutations observed in a validation set of patients that were not part of the eSTR discovery group (Methods). While one eSTR was mutated in 12 patients, most eSTRs were either mutated in zero (627) or one (269) patient(s). This meant statistical tests were underpowered to validate individual eSTRs. Instead, we determined the overall performance of our eSTR panel in predicting gene expression changes. We did this by comparing the observed gene expression change (increase or decrease) between healthy and tumour samples with the change predicted by the *β* of each mutated eSTR. For 67.1% of eSTR mutations, gene expression changed in the direction predicted by *β* . This was significantly better than randomly guessing whether gene expression would increase or decrease with probability 0.5 (Binomial test, *P*-value *≪* 0.05).

Next, we calculated an expected mutation impact for each eSTR mutation, which we defined as the product of the absolute value of *β* (*β_abs_*) and the difference in allele length (Δ*_l_*). *β_abs_* is a measure of how much gene expression is expected to change if the mean allele length were to change by one unit. Thus, by multiplying *β_abs_* with Δ*_l_*, we get an estimate of how much a particular eSTR mutation is expected to change gene expression between the healthy and tumour tissue of a patient. We found that mutations with higher expected impact lead to better predictions of gene expression changes (Figure 3D). Using only the quartile of mutations with the highest expected impact, the proportion of correctly predicted expression changes rose to 75.9% (283 / 373 mutations). This was higher than the accuracies obtained using the first, second, or third quartiles of expected mutation impact, which were 56.4%, 64.9%, and 71.3%, respectively.

To test whether eSTRs generated from non-cancer sequencing samples could predict gene expression changes following somatic mutations, we repeated our validation approach with the eSTR panel presented in Fotsing *et al*. (2019). We converted the Fotsing panel to GRCh38 coordinates using LiftOver (Hinrichs *et al*., 2006) and intersected it with our STR panel using BEDTools 2.30.0 (Quinlan and Hall, 2010). Fotsing *et al*. (2019) generated their eSTR panel from whole-genome sequencing (WGS) data, and STRs in a 100kb window around genes were considered as potential eSTRs. This meant that only a small number of Fotsing eSTRs could be mapped to STRs included in our WES-based analysis. In our validation set, we observed 429 mutations across 155 Fotsing eSTR loci. Only four of these were fine-mapped eSTRs that Fotsing *et al*. (2019) estimated to be causal for gene expression regulation. It is, therefore, perhaps not surprising that predictions of gene expression changes for this set of mutations were not significantly different from random guessing (Binomial test, *P*-value 0.87).

### eSTRs are more mutable in MSI tumours

Since we could associate eSTR mutations with gene expression changes in CRC tumours, we wondered whether tumours make use these effects to reprogram their phenotype. One piece of evidence for this would be if eSTRs are more mutable compared to non-eSTRs. This could indicate a selective advantage for tumours where the gene expression profile is changed due to eSTR mutations. Thus, we compared the mutability of eSTRs and non-eSTRs for MSS and MSI patients (Methods). For this analysis, STRs were grouped into repeat types of the same unit size and allele length, since we found that these characteristics have a large influence on STR mutability (Figure 2). For MSS patients, eSTRs were more mutable than non-eSTRs for five out of 46 tested repeat types. This was not significantly different from a null distribution obtained under random permutations of eSTR labels (permutation test, *P*-value = 0.920) (Figure 4). For MSI patients, on the other hand, an increase in eSTR mutability was observed for 12 out of 27 tested repeat types. This was significantly higher than the corresponding null distribution (permutation test, *P*-value = 1.00e*^−^*^4^) (Figure 4). The fraction of repeat types for which non-eSTRs were more mutable than eSTRs was not significantly different from random expectation for MSS (permutation test, *P*-value = 0.17) or MSI patients (permutation test, *P*-value = 0.19) (Supplementary file 1: Figure S5).

**Figure 4.**
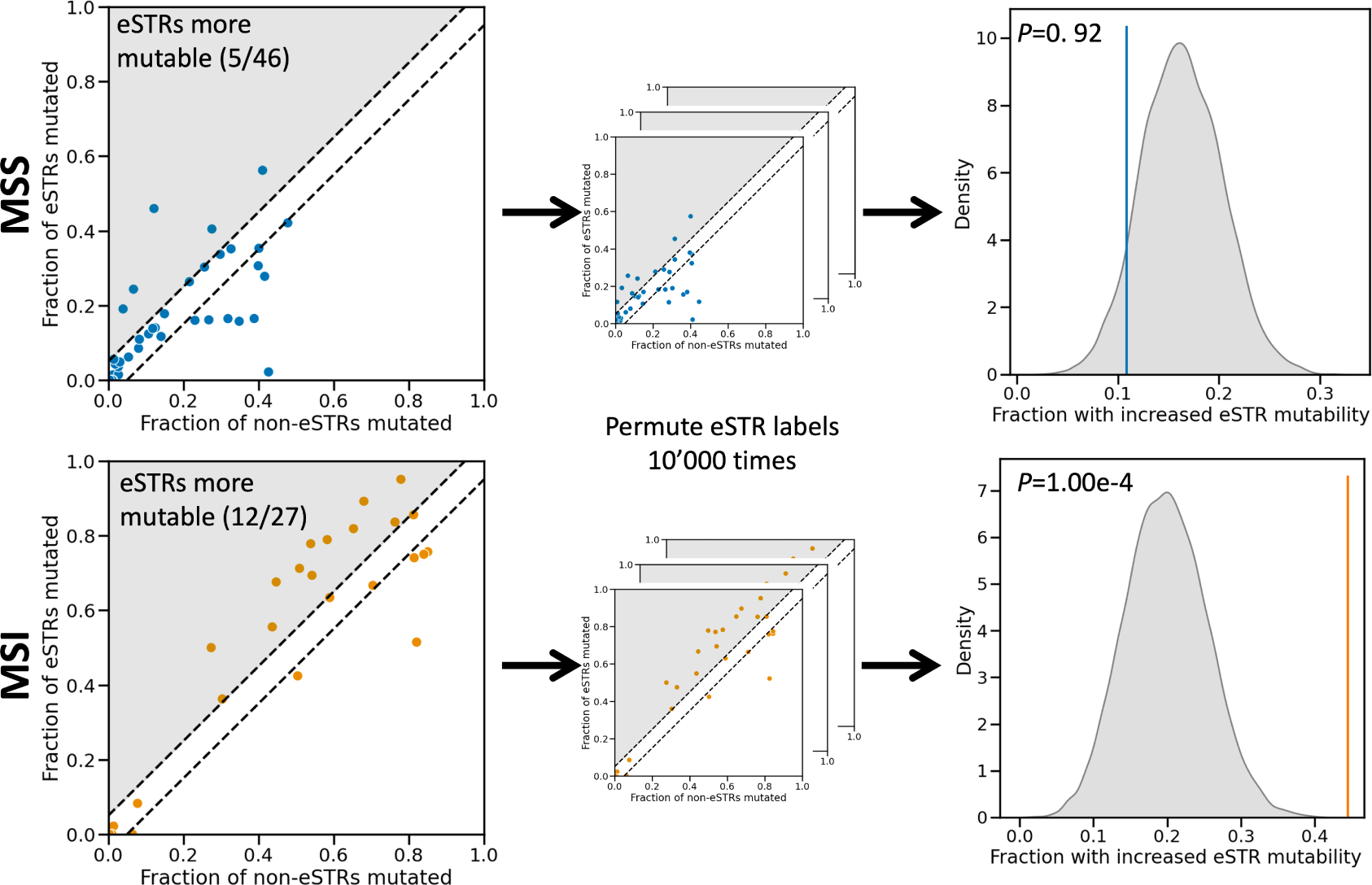
Comparing the mutability of eSTRs and non-eSTRs in CRC tumours. The top row shows results for MSS patients, the bottom row for MSI patients. In the scatter plots on the left every dot represents a repeat type, which is uniquely characterised by a combination of STR unit size and allele length. The fraction of mutated non-eSTRs is shown on the x-axis, and the fraction of mutated eSTRs on the y-axis. Dots that fall between the dashed lines represent repeat types for which no difference in mutability between eSTRs and non-eSTRs was observed. For repeat types that fall in the shaded region, eSTRs were more mutable than non-eSTRs (their numbers are noted in the top left). To generate a null distribution, eSTR labels were permuted 10000 times for both MSS and MSI patient mutation data (middle column). For each permutation, the fraction of repeat types for which the eSTRs were more mutable was determined. Kernel density estimates of the resulting distributions are shown in the right column. Vertical coloured stripes represent the observed fraction of repeat types where eSTRs were more mutable. *P*-values obtained from comparing the observed values to their respective null distributions using permutation tests are shown in the top left. Abbreviations: MSS, microsatellite stable; MSI, microsatellite instable; STR, expression short tandem repeat.

## DISCUSSION

Allele length variations at STR loci are known to regulate gene expression in healthy tissue (Gymrek *et al*. (2016); Fotsing *et al*. (2019); Shi *et al*. (2023)). While STRs are a rich source of somatic mutations in CRC — particularly in tumours with the MSI phenotype — it is still unclear if and how these mutations influence cancer phenotypes by modulating gene expression in tumours. Here, we describe for the first time a set of eSTRs for which the allele length is associated with gene expression in CRC tumours. Furthermore, we demonstrate that the linear models underlying the eSTR-expression associations allow for predictions of gene expression changes in response to somatic mutations in tumours. We also show evidence for an increased mutability of eSTRs compared to similar non-eSTR loci in MSI, but not MSS tumours. This could point to a selective advantage for eSTR mutations that affect the gene expression profile in MSI tumours.

First, we generated a novel STR annotation of the human protein-coding genome, which we genotyped in WES samples from TCGA. We identified somatic STR mutations by comparing STR allele lengths between patient-matched tumour and healthy samples. As expected, STRs were more frequently mutated in MSI tumours than in MSS tumours. Interestingly, we only observed an increase in STR deletions in MSI tumours, with no significant difference in insertion frequencies between MSI and MSS tumours. While a bias towards deletions for STR mutations in MSI tumours has been reported previously (Maruvka *et al*., 2017), our data indicates that this may be an understatement: the MSI phenotype seems to selectively increase STR deletions, without significantly affecting insertion frequencies. This selective increase in STR deletions in MSI tumours has been observed as far back as 1993 (Ionov *et al*., 1993), which we here confirm at a much larger scale in high-throughput data. For MSS tumours, the number of insertions and deletions was balanced. This is also inconsistent with Maruvka *et al*. (2017), where MSS tumours were reported to have more STR insertions than deletions. It is possible that differences in STR genotyping approach are responsible for this. Many STR genotyping tools are limited to calling STRs with short allele lengths from short-read sequencing data. Since STRs with short allele lengths are more prone to insertions (Xu *et al*., 2000), this might cause the appearance of an overall bias toward insertions for STRs in MSS samples. We used GangSTR to call STR allele lengths (Mitra *et al*., 2021). GangSTR is a dedicated STR genotyping algorithm that is less restricted by read length for determining STR allele lengths than most other tools. Since this is expected to improve the sampling of STRs with longer alleles — which are more prone to deletions (Xu *et al*., 2000) — this may lead to the symmetric distribution of insertions and deletions observed here. This is also in line with the fact that STR loci do not expand indefinitely and rarely disappear completely from the genome. STR variability rather tends to result in a stationary distribution around a central allele length (Xu *et al*. (2000); Sun *et al*. (2012)). Dedicated benchmarking experiments are needed to determine whether the inconsistencies between our findings and those of others are due to differences in STR mutation calling approaches.

Apart from the effect of MSI status, locus-specific characteristics also had an effect on STR mutability. STRs with longer unit sizes were less mutable than STRs with shorter unit sizes, whereas long allele lengths were associated with more frequent mutations. These findings are in line with previous studies of STR variability (Willems *et al*. (2014); Maruvka *et al*. (2017)). We observed that the MSI phenotype primarily affects mononucleotide repeats, which were over six times more mutable in MSI tumours compared to MSS tumours. While there was also a significant increase in the proportion of mutated di-, tri- and tertranucleotide repeats in MSI tumours, the relative differences to MSS tumours were smaller.

For STRs with unit sizes five and six no significant difference in mutability was observed between the phenotypes. This could mean that STRs of unit size five and six are not affected by the MSI phenotype. Alternatively, it may be because these classes of STR are much less abundant in the genome, thus constituting a smaller sample.

We present a novel panel of 1244 eSTRs that were associated with gene expression in colorectal cancer tumours. For 73 eSTRs, the allele length was related to the expression of a COSMIC cancer gene (Tate *et al*., 2019). However, a significant association does not guarantee that a putative eSTR is causal for gene expression changes: specific STR allele lengths may be in linkage disequilibrium with some untracked eQTL. Previous studies addressed this using statistical fine-mapping approaches where eSTR effects are disentangled from their genomic contexts (Gymrek *et al*. (2016); Fotsing *et al*. (2019)). Since our analyses were based on WES data, large parts of the genomes we investigated were not observable by us. This prevented us from accounting for any tagged variants using the statistical approaches described in other works. Instead, we quantified the ability of our putative eSTRs to predict gene expression changes in response to somatic mutations in CRC tumours. Our reasoning was that — if a putative eSTR indeed regulates gene expression — a somatic mutation at an eSTR should result in a difference in expression between the healthy and tumour sample. Furthermore, the direction of expression change should correspond to the sign of the slope of the linear model for that specific eSTR-gene pair. It would be highly unlikely for a tagged eQTL to mutate simultaneously and consistently with the eSTR across different patients. This approach should therefore be able to validate the causal nature of the relationship between gene expression and eSTR allele lengths. Furthermore, since this validation approach is based on comparisons between patient-matched samples, many biological confounding factors are accounted for.

Using the eSTR panel, we could predict the direction of gene expression changes in response to eSTR mutations with a significant degree of accuracy. Moreover, when focusing on eSTR mutations where we expected the largest impact on gene expression — and thus a stronger signal — our prediction accuracy increased. Using the quartile of eSTR mutations with the highest expected impact, we reached over 75% accuracy in predicting the direction of gene expression change. This fits with the idea that eSTRs can function as ’tuning knobs’ in gene regulation (Verbiest *et al*., 2023), with larger changes in allele lengths leading to bigger, more readily detectable shifts in gene expression.

Notably, a recent survey of STR mutations in cancer also attempted to identify gene regulatory effects of STR mutations (Fujimoto *et al*., 2020). While the authors investigated STR mutations across a cohort of 21 cancer types, they reported not a single significant association to gene expression. It is possible that all eSTR effects in cancer are highly tissue specific and therefore obscured in the pan-cancer analysis presented in Fujimoto *et al*. (2020). However, it seems more likely that methodological differences underlie the discrepancies between our findings. Fujimoto *et al*. (2020) defined two groups of tumour samples in their analysis: those with and without STR mutations in their promoter or UTR regions. They then tested for differences in gene expression levels between these two groups of tumours, without considering expression levels in patient-matched healthy samples. Instead, we reasoned that the effects of eSTR mutations should be observable by comparing gene expression levels between healthy and tumour tissues of patients with somatic eSTR mutations.

Finally, we found that eSTRs were mutated more often than non-eSTRs of the same unit size and allele length in MSI tumours. This might be an early indication that tumour subclones can gain a selective advantage by modifying their phenotype through eSTRs mutations. We did not observe an increase in eSTR mutability for MSS tumours. This could be related to the overall lower mutability of STRs in MSS tumours. In the absence of STR hypermutability MSS tumours may have to rely on other mechanisms to change their expression profiles. For example, it has been found that copy number variants (CNVs) are more common in MSS tumours compared to MSI tumours (The Cancer Genome Atlas Network, 2012). We observed this association when filtering out STR length calls that overlapped CNV events (Methods). In MSS patients, 17.1% of STR length calls overlapped a CNV event, compared to only 1.3% of STR calls in MSI patients.

Even though we found evidence of STR-mediated gene regulation in CRC, there are some limitations to the analyses presented here. First, our focus on WES data limited the genomic regions in which we could detect eSTRs. While previous studies have reported that eSTRs are enriched around transcription start sites, there are also eSTRs that are located far away from protein-coding genes (Fotsing *et al*., 2019). Our eSTR panel therefore likely represents a subset of STR loci that regulate gene expression in CRC. Second, with 16 patients our validation set was relatively small. This was because we required patients for which both WES and gene expression were available for both a primary tumour sample and a solid tissue normal sample. While we identified a sufficient number of eSTR mutations in this validation set to assess our eSTR panel as a whole, it did not allow for significance testing at the level of individual eSTR loci. This is also why we have mostly refrained from functionally interpreting specific eSTR loci or the genes they are associated with.

In a broader context, our results may be related to previous observations regarding gene expression profiles in MSI tumours. The consensus molecular subtypes (CMSs) are a gene expression-based classification system of CRC tumours (Guinney *et al*., 2015). Most MSI tumours belong to CMS1. In fact, MSI is one of the defining characteristics of CMS1 according to Guinney *et al*. (2015). Given our findings, it is possible that the hypermutability of eSTRs contributes to shaping a distinct gene expression landscape in MSI tumours. As noted above, however, there are other mutational mechanisms that have a non-random association to the MSI phenotype (The Cancer Genome Atlas Network, 2012). In our analyses, we filtered out STRs and genes that overlap somatic CNVs to prevent the effects of such variants from confounding our eSTR detection. Future studiesshould integrate data on CNVs, STRs, and other types of (structural) variants with expression changes in CRC tumours. This could elucidate the relative contributions of different mutational processes to the dysregulation of gene expression in MSI tumours.

## CONCLUSIONS

To the best of our knowledge, our findings represent the first cancer-based eSTR panel generated from high-throughput sequencing data. The alteration of eSTR repeat numbers is a largely unexplored way through which tumours can reprogram their phenotypes, and our results underscore the need for more research on this topic. Obvious next steps would be to extend our analyses to larger datasets and other cancer types, as well as widening the scope from exome- to genome scale. More data will allow for statistical validation of individual eSTR loci. Studying whether the eSTR associations reported here can be replicated in other cancer types could uncover whether eSTR associations are tissue-specific, or if they represent general regulatory elements in cancer. It is our hope that such investigations will deepen our understanding of the role STRs play in cancer beyond the tumour-level MSS/MSI label that has been in place for decades.

## AVAILABILITY OF DATA AND MATERIALS

GRCh38 reference coordinates for STRs genotyped in the analyses presented here are available for download from the WebSTR database: http://webstr.ucsd.edu/ (Lundström *et al*., 2023). The full, unfiltered STR panel is available from the corresponding authors on reasonable request. Notebooks containing the analyses presented here are available at https://github.com/acg-team/ `STRs-in-CRC`. This repository also contains dummy data that can be used by researchers who do not have access to restricted TCGA data to test and validate the implementation of analyses presented here. All cancer sequencing data were generated by the TCGA Research Network: https://www.cancer.gov/tcga. Data from the COAD and READ cohorts were downloaded from the GDC knowledge base (data release version 31.0). Access to restricted TCGA data was granted under dbGaP study phs000178.v11.p8.c1.

## AUTHORS’ CONTRIBUTIONS

MAV designed and performed the analyses, interpreted data, developed software, and wrote the manuscript. OL acquired data, performed analyses, and developed software. FX performed analyses. MB and TBS contributed to discussions, the interpretation of data, and the writing and revising of the manuscript. MA acquired the funding and contributed to the conception of the study, the interpretation of data, and the writing and revising of the manuscript. All authors read and approved the final manuscript.

## CONFLICTS OF INTEREST

The authors declare that they have no conflicts of interest.

## ACKNOWLEDGEMENTS

MAV, OL, FX, and MA were supported by SNSF Sinergia grant CRSII5 193832 and the EU Horizon 2020 research and innovation program under the Marie Skłodowska-Curie grant agreement No. 823886. TBS and MA acknowledge that they received the ”Scientific Exchanges” SNSF grant IZSEZ0 203264. The authors acknowledge Andrey Ershov for adding an interactive graph illustrating the results of this paper to the WebSTR web interface. Furthermore, the authors would like to express their gratitude to all specimen donors who provided samples to the TCGA COAD and READ cohorts.

## Notes

### Competing Interest Statement

The authors have declared no competing interest.

## REFERENCES

1. Avvaru, A. K., Sowpati, D. T., and Mishra, R. K. (2018). PERF: an exhaustive algorithm for ultra-fast and efficient identification of microsatellites from large DNA sequences. *Bioinformatics (Oxford*, England), 34(6):943–948.

2. Benson, G. (1999). Tandem repeats finder: a program to analyze DNA sequences. Nucleic Acids Research, 27(2):573–580.

3. Bilgin Sonay, T., Koletou, M., and Wagner, A. (2015). A survey of tandem repeat instabilities and associated gene expression changes in 35 colorectal cancers. BMC Genomics, 16(1):702.

4. Boland, C. R. and Goel, A. (2010). Microsatellite Instability in Colorectal Cancer. Gastroenterology, 138(6):2073–2087.e3.

5. Bonneville, R., Krook, M. A., Kautto, E. A., Miya, J., Wing, M. R., Chen, H.-Z., Reeser, J. W., Yu, L., and Roychowdhury, S. (2017). Landscape of Microsatellite Instability Across 39 Cancer Types. JCO Precision Oncology, (1):1–15.

6. Delucchi, M., Näf, P., Bliven, S., and Anisimova, M. (2021). TRAL 2.0: Tandem Repeat Detection With Circular Profile Hidden Markov Models and Evolutionary Aligner. Frontiers in Bioinformatics, 1.

7. Eddy, S. R. (2011). Accelerated Profile HMM Searches. PLOS Computational Biology, 7(10):e1002195.

8. Ellegren, H. (2004). Microsatellites: Simple sequences with complex evolution. Nature Reviews Genetics, 5(6):435–445.

9. Fotsing, S. F., Margoliash, J., Wang, C., Saini, S., Yanicky, R., Shleizer-Burko, S., Goren, A., and Gymrek, M. (2019). The impact of short tandem repeat variation on gene expression. Nature Genetics, 51(11):1652–1659.

10. Fujimoto, A., Fujita, M., Hasegawa, T., Wong, J. H., Maejima, K., Oku-Sasaki, A., Nakano, K., Shiraishi, Y., Miyano, S., Yamamoto, G., Akagi, K., Imoto, S., and Nakagawa, H. (2020). Comprehensive analysis of indels in whole-genome microsatellite regions and microsatellite instability across 21 cancer types. Genome Research, 30(3):334–346.

11. Guinney, J., Dienstmann, R., Wang, X., De Reyniès, A., Schlicker, A., Soneson, C., Marisa, L., Roepman, P., Nyamundanda, G., Angelino, P., Bot, B. M., Morris, J. S., Simon, I. M., Gerster, S., Fessler, E., De Sousa .E Melo, F., Missiaglia, E., Ramay, H., Barras, D., Homicsko, K., Maru, D., Manyam, G. C., Broom, B., Boige, V., Perez-Villamil, B., Laderas, T., Salazar, R., Gray, J. W., Hanahan, D., Tabernero, J., Bernards, R., Friend, S. H., Laurent-Puig, P., Medema, J. P., Sadanandam, A., Wessels, L., Delorenzi, M., Kopetz, S., Vermeulen, L., and Tejpar, S. (2015). The consensus molecular subtypes of colorectal cancer. Nature Medicine, 21(11):1350–1356.

12. Gymrek, M., Willems, T., Guilmatre, A., Zeng, H., Markus, B., Georgiev, S., Daly, M. J., Price, A. L., Pritchard, J. K., Sharp, A. J., and Erlich, Y. (2016). Abundant contribution of short tandem repeats to gene expression variation in humans. Nature Genetics, 48(1):22–29.

13. Hause, R. J., Pritchard, C. C., Shendure, J., and Salipante, S. J. (2016). Classification and characterization of microsatellite instability across 18 cancer types. Nature Medicine, 22(11):1342–1350.

14. Hinrichs, A. S., Karolchik, D., Baertsch, R., Barber, G. P., Bejerano, G., Clawson, H., Diekhans, M., Furey, T. S., Harte, R. A., Hsu, F., Hillman-Jackson, J., Kuhn, R. M., Pedersen, J. S., Pohl, A., Raney, B. J., Rosenbloom, K. R., Siepel, A., Smith, K. E., Sugnet, C. W., Sultan-Qurraie, A., Thomas, D. J., Trumbower, H., Weber, R. J., Weirauch, M., Zweig, A. S., Haussler, D., and Kent, W. J. (2006). The UCSC Genome Browser Database: update 2006. Nucleic Acids Research, 34(Database issue):D590– D598.

15. Huang, Q., Carrio-Cordo, P., Gao, B., Paloots, R., and Baudis, M. (2021). The Progenetix oncogenomic resource in 2021. Database, 2021:baab043.

16. Ionov, Y., Peinado, M. A., Malkhosyan, S., Shibata, D., and Perucho, M. (1993). Ubiquitous somatic mutations in simple repeated sequences reveal a new mechanism for colonic carcinogenesis. Nature, 363(6429):558–561.

17. Kim, T.-M., Laird, P. W., and Park, P. J. (2013). The Landscape of Microsatellite Instability in Colorectal and Endometrial Cancer Genomes. Cell, 155(4):858–868.

18. Lai, Y. and Sun, F. (2003). The Relationship Between Microsatellite Slippage Mutation Rate and the Number of Repeat Units. Molecular Biology and Evolution, 20(12):2123–2131.

19. Lundström, O. S., Verbiest, M. A., Xia, F., Jam, H. Z., Zlobec, I., Anisimova, M., and Gymrek, M. (2023). WebSTR: A Population-wide Database of Short Tandem Repeat Variation in Humans. Journal of Molecular Biology, page 168260.

20. Martin-Trujillo, A., Garg, P., Patel, N., Jadhav, B., and Sharp, A. J. (2023). Genome-wide evaluation of the effect of short tandem repeat variation on local DNA methylation. Genome Research, 33(2):184–196.

21. Maruvka, Y. E., Mouw, K. W., Karlic, R., Parasuraman, P., Kamburov, A., Polak, P., Haradhvala, N. J., Hess, J. M., Rheinbay, E., Brody, Y., Koren, A., Braunstein, L. Z., D’Andrea, A., Lawrence, M. S., Bass, A., Bernards, A., Michor, F., and Getz, G. (2017). Analysis of somatic microsatellite indels identifies driver events in human tumors. Nature Biotechnology, 35(10):951–959.

22. Mayer, C., Leese, F., and Tollrian, R. (2010). Genome-wide analysis of tandem repeats in Daphnia pulex - a comparative approach. BMC Genomics, 11(1):277.

23. Mitra, I., Huang, B., Mousavi, N., Ma, N., Lamkin, M., Yanicky, R., Shleizer-Burko, S., Lohmueller, K. E., and Gymrek, M. (2021). Patterns of de novo tandem repeat mutations and their role in autism. Nature, 589(7841):246–250.

24. Mousavi, N., Margoliash, J., Pusarla, N., Saini, S., Yanicky, R., and Gymrek, M. (2021). TRTools: a toolkit for genome-wide analysis of tandem repeats. Bioinformatics, 37(5):731–733.

25. Mousavi, N., Shleizer-Burko, S., Yanicky, R., and Gymrek, M. (2019). Profiling the genome-wide landscape of tandem repeat expansions. Nucleic Acids Research, 47(15):e90–e90.

26. Newman, A. M. and Cooper, J. B. (2007). XSTREAM: A practical algorithm for identification and architecture modeling of tandem repeats in protein sequences. BMC Bioinformatics, 8(1):382.

27. Quinlan, A. R. and Hall, I. M. (2010). BEDTools: a flexible suite of utilities for comparing genomic features. Bioinformatics, 26(6):841–842.

28. Schaper, E., Korsunsky, A., Pečerska, J., Messina, A., Murri, R., Stockinger, H., Zoller, S., Xenarios, I., and Anisimova, M. (2015). TRAL: tandem repeat annotation library. Bioinformatics, 31(18):3051– 3053.

29. Seabold, S. and Perktold, J. (2010). Statsmodels: Econometric and Statistical Modeling with Python. In Walt, S. v. d. and Millman, J., editors, Proceedings of the 9th Python in Science Conference, pages 92 – 96.

30. Shi, Y., Niu, Y., Zhang, P., Luo, H., Liu, S., Zhang, S., Wang, J., Li, Y., Liu, X., Song, T., Xu, T., and He, S. (2023). Characterization of genome-wide STR variation in 6487 human genomes. Nature Communications, 14(1):2092.

31. Sun, J. X., Helgason, A., Masson, G., Ebenesersdóttir, S. S., Li, H., Mallick, S., Gnerre, S., Patterson, N., Kong, A., Reich, D., and Stefansson, K. (2012). A direct characterization of human mutation based on microsatellites. Nature Genetics, 44(10):1161–1165.

32. Tate, J. G., Bamford, S., Jubb, H. C., Sondka, Z., Beare, D. M., Bindal, N., Boutselakis, H., Cole, C. G., Creatore, C., Dawson, E., Fish, P., Harsha, B., Hathaway, C., Jupe, S. C., Kok, C. Y., Noble, K., Ponting, L., Ramshaw, C. C., Rye, C. E., Speedy, H. E., Stefancsik, R., Thompson, S. L., Wang, S., Ward, S., Campbell, P. J., and Forbes, S. A. (2019). COSMIC: the Catalogue Of Somatic Mutations In Cancer. Nucleic Acids Research, 47(D1):D941–D947.

33. The Cancer Genome Atlas Network (2012). Comprehensive molecular characterization of human colon and rectal cancer. Nature, 487(7407):330–337.

34. Verbiest, M., Maksimov, M., Jin, Y., Anisimova, M., Gymrek, M., and Bilgin Sonay, T. (2023). Mutation and selection processes regulating short tandem repeats give rise to genetic and phenotypic diversity across species. Journal of Evolutionary Biology, 36(2):321–336.

35. Willems, T., Gymrek, M., Highnam, G., Mittelman, D., and Erlich, Y. (2014). The landscape of human STR variation. Genome Research, 24(11):1894–1904.

36. Xu, X., Peng, M., Fang, Z., and Xu, X. (2000). The direction of microsatellite mutations is dependent upon allele length. Nature Genetics, 24(4):396–399.

37. Zhao, H. and Baudis, M. (2023). labelSeg: segment annotation for tumor copy number alteration profiles. Preprint at bioRxiv 10.1101/2023.05.17.541097.

