## Supplementary File 1 for "Short tandem repeat mutations regulate gene expression in colorectal cancer"

Supplementary figures for:

**Short tandem repeat mutations regulate**

**gene expression in colorectal cancer**

Verbiest *et al*.

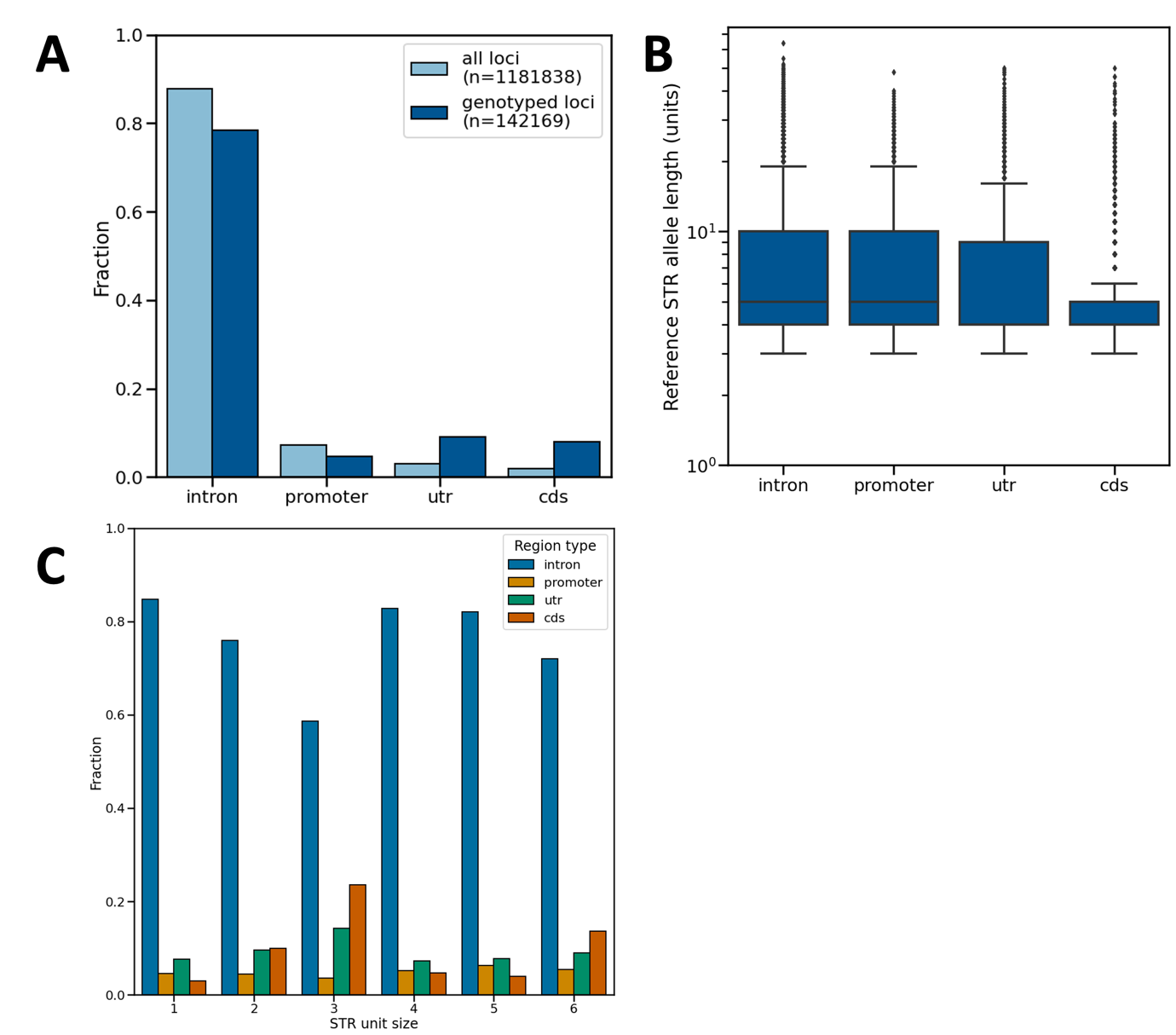

**Figure S1** A novel STR panel for human protein-coding genes. A) Distribution of STR loci across genomic region types. The distribution of the full STR panel, as well as the subset of STRs for which allele lengths could be called from TCGA WES samples are shown. B) Comparison of reference allele lengths across different genomic region types for genotyped STRs. Boxes extend from Q1 to Q3, with a line indicating the median value. C) Genomic locations of STRs with different unit sizes. Abbreviations: UTR, untranslated region; CDS, coding sequence.

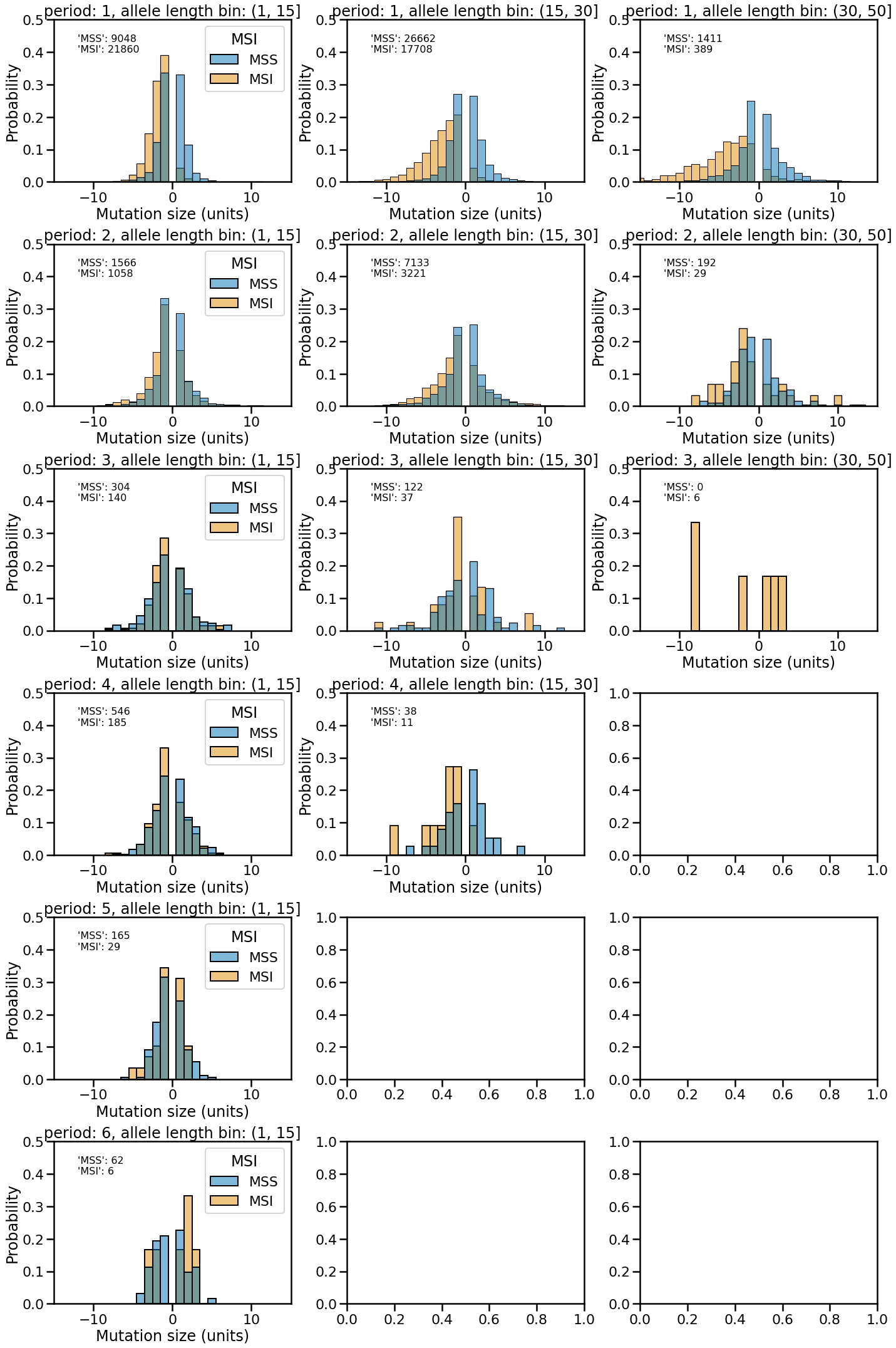

**Figure S2** Distributions of STR mutation step sizes (in units) for MSS and MSI patients. Each row of panels shows data from one unit size (one through six), and each column shows data from a range of allele lengths ((1, 15], (15, 30], (30, 50]). Negative step sizes indicate deletions in tumours and positive step sizes insertions. The y-axis displays the probability of an STR mutation being a certain step size. Data from MSS (blue) and MSI (orange) tumours are shown separately as overlapping histograms. The histograms for MSS and MSI data each sum to one. Abbreviations: MSS, microsatellite stable; MSI, microsatellite instable.

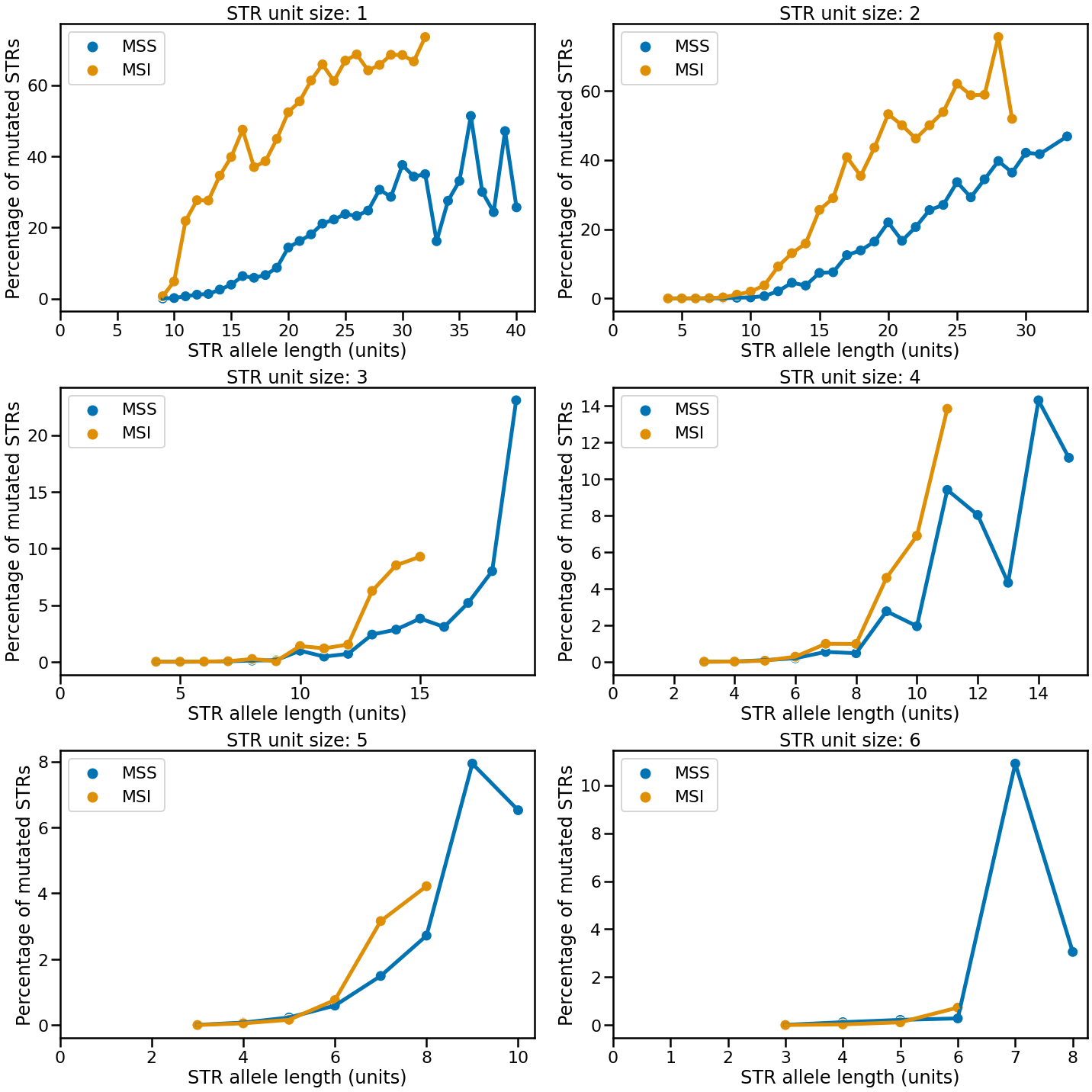

**Figure S3** STR mutation rates as a function of unit size and allele length. For each unit size, mutation frequencies are shown for STR allele lengths for which a comparison between a healthy and tumour sample could be made in at least 50 patients. Results are plotted separately for MSS and MSI tumours. Abbreviations: MSS, microsatellite stable; MSI, microsatellite instable.

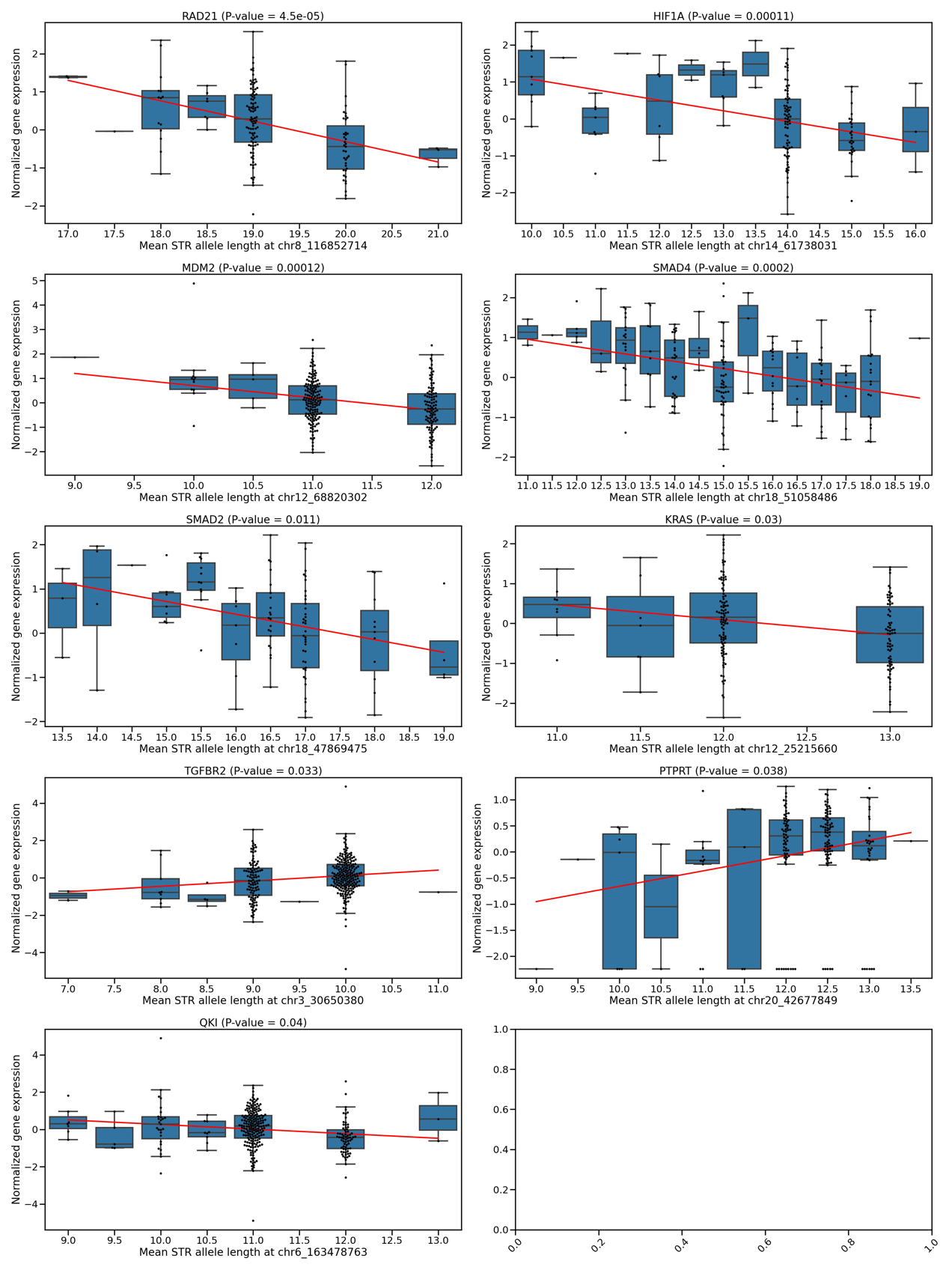

**Figure S4** eSTRs associated with expression of CRC-related genes. The STR length is shown on the x-axes (mean of two alleles), and the normalised gene expression on the y-axes. Every dot represents one tumour sample. Boxplots show the distribution of expression values across tumours at each STR genotype. Boxes extend from Q1 to Q3, with a line indicating the median value. The red line represents the linear model relating STR length to normalised expression.

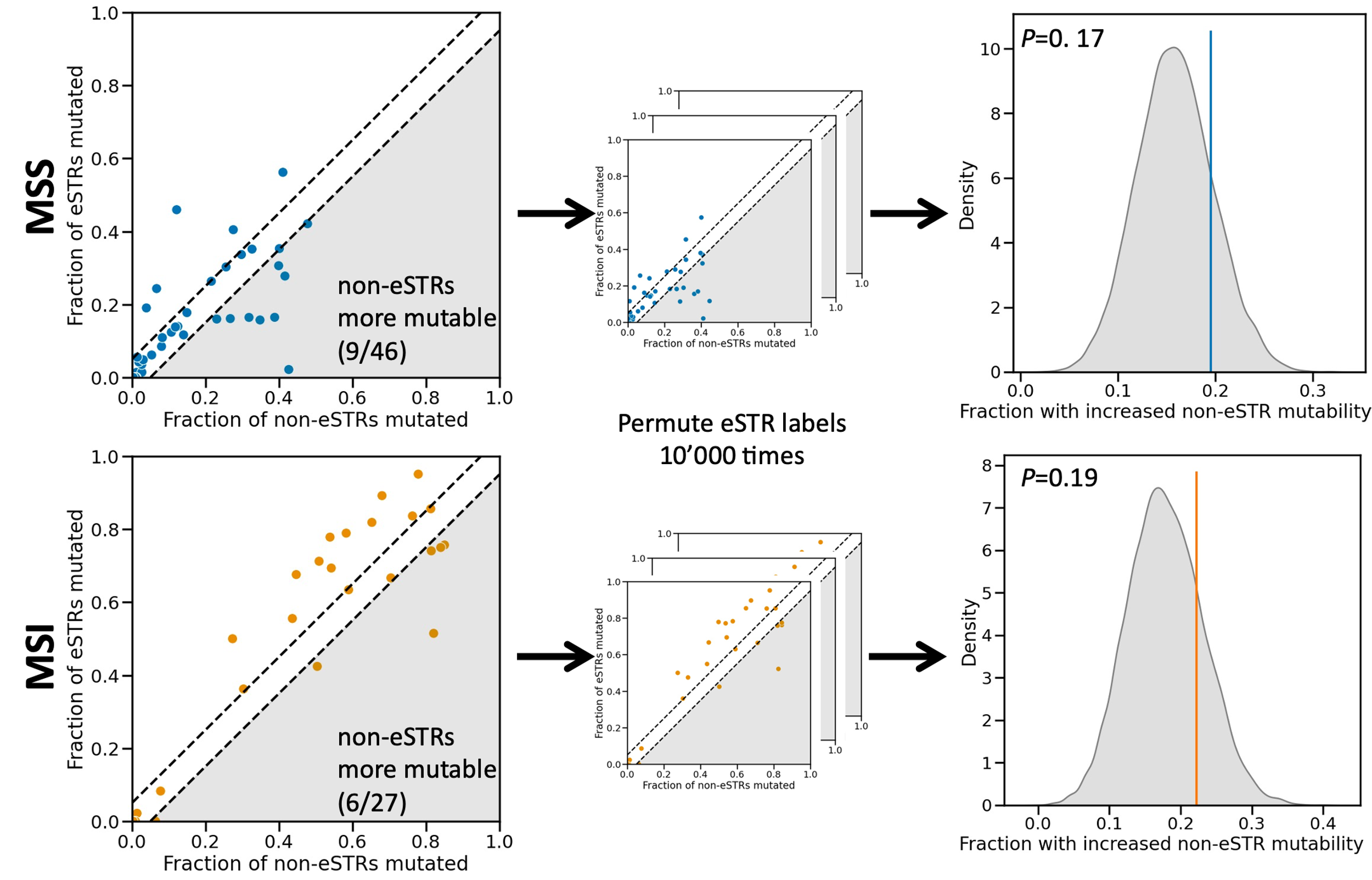

**Figure S5** Comparing the mutability of eSTRs and non-eSTRs in CRC tumours. The top row represents results obtained for MSS patients, the bottom row for MSI patients. In the scatter plots on the left every dot represents a repeat type, which is uniquely characterised by a combination of STR unit size and allele length. The fraction of mutated non-eSTRs is shown on the x-axis, and the fraction of mutated eSTRs on the y-axis. Dots that fall between the dashed lines represent repeat types for which no difference in mutability between eSTRs and non-eSTRs was observed. For repeat types that fall in the shaded region, non-eSTRs were more mutable than eSTRs (their numbers are noted in the bottom right). To generate a null distribution, eSTR labels were permuted 10000 times for both MSS and MSI patient mutation data (middle column). For each permutation, the fraction of repeat types for which the non-eSTRs were more mutable was determined. Kernel density estimates of the resulting distributions are shown in the right column. Vertical coloured stripes represent the observed fraction of repeat types where non-eSTRs were more mutable. *P*-values obtained from comparing the observed values to their respective null distributions using permutation tests are shown in the top left. Abbreviations: MSS, microsatellite stable; MSI, microsatellite instable; STR, expression short tandem repeat.
